# A novel protistan trait database reveals functional redundancy and complementarity in terrestrial protists (Amoebozoa & Rhizaria)

**DOI:** 10.1101/2025.03.28.645897

**Authors:** Jule Freudenthal, Martin Schlegel, Michael Bonkowski, Kenneth Dumack

**Affiliations:** Terrestrial Ecology, Institute of Zoology, Cluster of Excellence on Plant Sciences (CEPLAS), University of Cologne, Zülpicher Str. 47b, 50674 Köln, Germany; Biodiversity and Evolution, Institute of Biology, University Leipzig, Talstraße 33, 04103 Leipzig, Germany; German Centre for Integrative Biodiversity Research (iDiv) Halle Jena Leipzig, Puschstraße 4, 04103 Leipzig, Germany

## Abstract

The inclusion of functional traits of protists in environmental sequencing surveys, in addition to the traditional taxonomic framework, is essential for a better understanding of their roles and impacts on ecosystem processes. We provide a database of functional traits for a widespread and important clade of protists, the Amoebozoa, based on extensive literature research in eight trait categories: Habitat, locomotion, nutrition, morphology, morphotype, size, spore formation, and disease-relatedness. The comparison of community traits of the Amoebozoa with sympatric but highly divergent Cercozoa (Rhizaria) revealed both convergent evolution of morphology or locomotion and distinct differences in habitat preference and feeding selectively. Amoebozoa seem to be rather unselective in their prey choice compared to Cercozoa. Indeed, the feeding preferences of Amoebozoa appeared to be related to cell size, whereas Cercozoa selectively feed on prey. Applications to metatranscriptomic data from soil, litter, and bark surfaces revealed differences in the average community trait compositions and ecosystem functioning, such as an increased proportion of disease-related Amoebozoa in soil or different proportions of nutrition types of Amoebozoa and Cercozoa on bark. This database will facilitate ecological analyses of sequencing data and improve our understanding of the diversity of adaptations of Amoebozoa to the environment and their functional roles in ecosystems.

## 1. Introduction

Refined information on the functional diversity of organisms, in addition to the traditional taxonomic framework, may greatly improve our knowledge of their function in ecosystem processes (Bouskill et al., 2012; Krause et al., 2014), but also, for example, how abiotic and biotic drivers shape communities (Briones, 2014; Fiore-Donno et al., 2019). To meet the analytical demands of environmental sequencing projects, trait-based data must be collated and tools developed to easily assign functional traits to existing sequencing databases.

Trait-based community analyses aim to link species diversity to ecosystem functioning (Lavorel and Garnier, 2002; Violle et al., 2007). Traits, on the one hand, determine the performance and fitness of an organism (response traits) by directly reflecting its adaptations to physical, chemical, and biotic environmental drivers. On the other hand, traits such as feeding mode capture their potential impact (effect traits) on the environment and, thus, species’ contributions to ecosystem functioning (Krause et al., 2014; Suding et al., 2008). Accordingly, trait-based community analyses may provide detailed information on the niche space occupied by communities (Lennon et al., 2012) or covered by a taxonomic group (Díaz et al., 2016) but also allow for an upscaling of ecosystem processes (Mulder et al., 2013).

Trait-based surveys are widely established for plants, animals, and prokaryotes (Beier et al., 2022; Bouskill et al., 2012; Louca et al., 2016). However, a sound functional understanding of the super-diverse communities of microbial eukaryotes is challenging, as their over 20 phyla comprise a multitude of completely independent evolutionary trajectories (Ruggiero et al., 2015). Furthermore, taxonomic and functional diversity are generally not necessarily coupled (Louca et al., 2016). For example, closely related taxa may exhibit different predatory impacts (Glücksman et al., 2010). Accordingly, studies covering a broad range of the diversity of microbial eukaryotes may so far provide only limited information on their functions (Aslani et al., 2022; Giachello et al., 2023; Köninger et al., 2023).

Over the past decade, there has been an increase in trait-based environmental surveys focusing on the lesser-investigated part of the microbial diversity, the protists (Amacker et al., 2022; Fiore-Donno et al., 2019; Flues et al., 2017; Jauss et al., 2021; Lamentowicz et al., 2020). The vast majority of microbial eukaryotic diversity is represented by protists, a paraphyletic assemblage of mostly unicellular eukaryotes. Protists fulfill numerous functions in terrestrial and aquatic ecosystems, like primary production and the exertion of distinct predation patterns, but some taxa are also important parasites of plants and animals. It is crucial to include functional traits to understand their roles in ecosystem processes. For example, the metabolic basis of protistan functional traits has been used to identify the main drivers of the shift between net heterotrophy and autotrophy in the oceans and to establish models predicting phytoplankton blooms (Alexander et al., 2015). Moreover, the importance of symbioses among the planktonic eukaryotes was only revealed after compiling the planktonic Protist Interaction DAtabase (PIDA, Bjorbækmo et al., 2020).

In soils, Amoebozoa and Cercozosa (Rhizaria) are the most dominant terrestrial protistan supergroups (Domonell et al., 2013; Dumack et al., 2016; Fiore-Donno et al., 2024; Urich et al., 2008; Voss et al., 2019). A detailed trait database exists for the Cercozoa and Endomyxa (Dumack et al., 2020). This trait database allows an easy assignment of traits to environmental sequences and thus enables functional insights into the structure of microbial food webs. Fiore-Donno et al. (2019) showed a relative increase in the abundance of shell-bearing Cercozoa with drier soils, supporting the long-assumed function of their shells, i.e., increased drought resistance due to reduced evaporation.

The supergroup Amoebozoa is equally diverse (Fig. 1) and abundant as Cercozoa and comprises three major lineages (Kang et al., 2017; Tekle et al., 2022): First, the Evosea, some of which with flagellated cells and complex life cycles, some are giant such as the plasmodia of Myxomycetes (clade: Cutosea and Conosea). Second, the Discosea, comprising the Flabellinia with flattened cells with separate hyaloplasm of which (lobose) subpseudopodia protrude (orders: Stygamoebida, Thecamoeboda, Dactylopodida, Vannellida, Dermamoebida), and the Centramoebida, some of which with scales (orders: Acanthamoebida, Pellitida, Himatismenida). Third, the Tubulinea, comprising shell-bearing but mostly naked lobose amoebae, with highly variable cell sizes ranging from 20 µm to several centimeters, taxa with larger cells often with branching (ramose) or network-forming (reticulose) pseudopodia (orders: Leptomyxida, Euamoebida, Arcellinida, Echinamoebida, Corycida). Moreover, the Amoebozoa include important human parasites like *Acanthamoeba* spp., *Balamuthia mandrillaris*, and *Entamoeba histolytica* (Fiore-Donno et al., 2016; Geisen et al., 2014; Tice et al., 2016; Walochnik, 2018). Traditional approaches early on identified the significance of Amoebozoa in soil systems (Azam et al., 1983; Clarholm, 1985; de Ruiter et al., 1995). Unfortunately, the widespread use of “general eukaryotic” primers in metabarcoding studies led to a consistent and dramatic underrepresentation of amoebozoan sequences in surveys of terrestrial eukaryote diversity. Metatranscriptomics does not suffer from these extreme primer biases and led to sequencing results largely concordant with microscopic surveys illustrating the dominance of Amoebozoa (Fiore-Donno et al., 2024; Freudenthal et al., 2022; Heck et al., 2023; Voss et al., 2019). Now, as the molecular methodology to assess the taxonomic richness and diversity of Amoebozoa is established, a trait database is highly needed to understand the diversity of their functional roles in terrestrial and aquatic communities.

**Figure 1:**
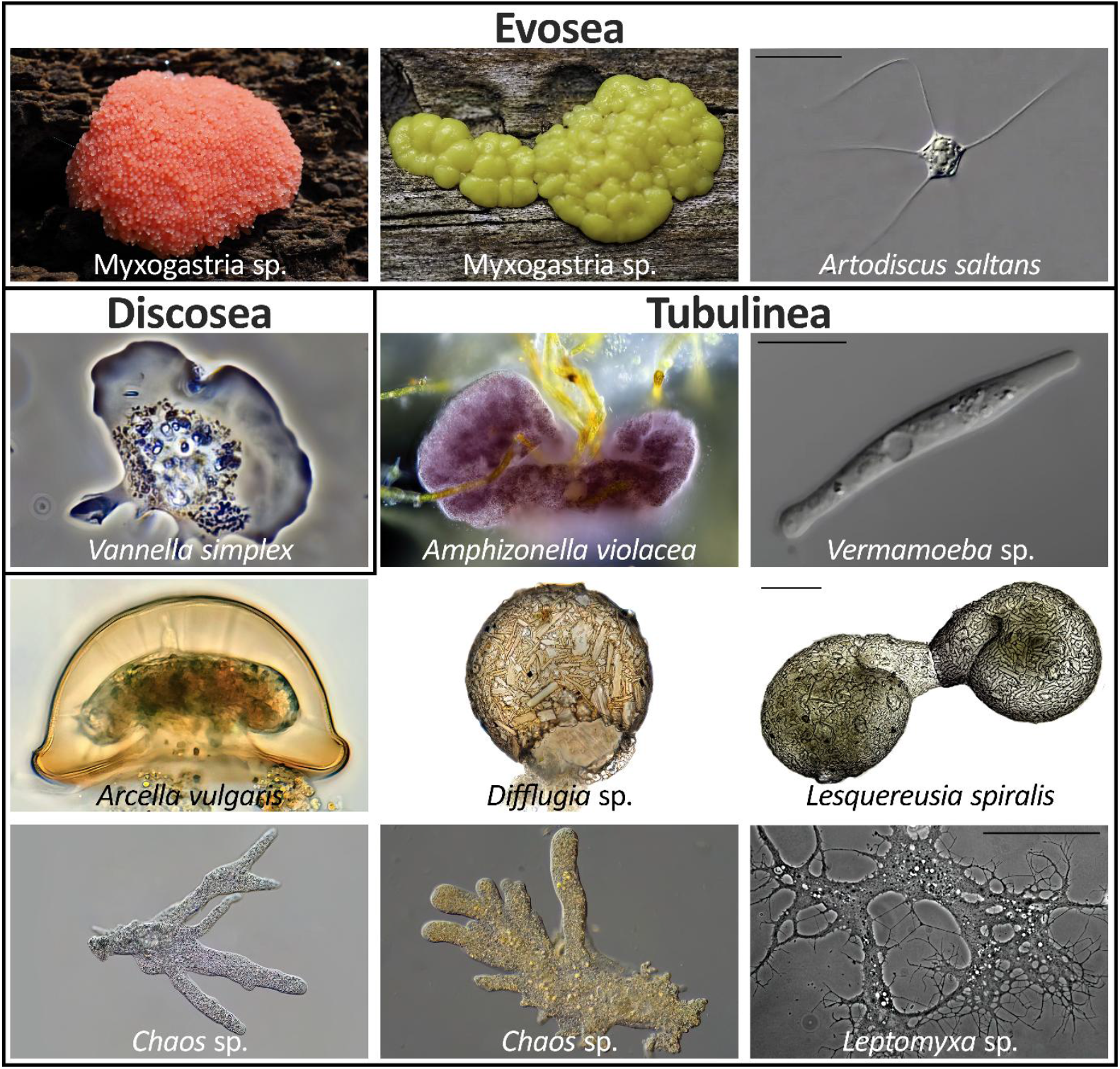
Overview of the morphological diversity of the Amobozoa. The graphic shows (1) the Evosea, represented by Myxomycetes sp. and *Artodiscus saltans* (Conosea), (2) the Discosea, represented by *Vannella simplex* (Flabellinia), and (3) the Tubulinea, represented by *Amphizonella violacea* (Corycida), *Vermamoeba* sp. (Echinamoebida), *Arcella vulgaris, Difflugia* sp., *Lesquereusia spiralis* (Arcellinida), *Chaos* sp. (Euamoebida), and *Leptomyxa* sp. (Leptomyxida).

Here, we provide a trait database for Amoebozoa to serve as a common reference and to facilitate functional ecological studies. We showcase the usage of amoebozoan traits on recently published metatranscriptomic data of soil, leaf litter, and bark surfaces, and we compare the traits of the two most dominant soil protistan supergroups – Amoebozoa and Cercozoa.

## 2. Materials and Methods

### 2.1 Intentional use of the trait database as a justification of its structure and content

As a baseline for our literature research, we screened the curated diversity of amoebozoan 18S rDNA sequences in the PR^2^ database v. 5.0.0 (Guillou et al., 2012) for all currently included amoebozoan genera. We attributed traits by means of the literature, including original descriptions, or accessing other available meta-analyses and already collated data. All consulted references are provided in the database. We attempted to be as exhaustive as possible in selecting functional traits. However, given the vast divergence within Amoebozoa, the trait database is still a strongly simplified representation of their functional diversity. Even though the database may not fulfill the expectations of taxonomists, it is primarily meant to facilitate functional analyses of sequencing data, especially by (microbial) ecologists who are not necessarily experts in protistan diversity. Accordingly, it was crucial to provide traits in discrete categories that can be easily subjected to statistical analyses, i.e., each trait for any taxon can only be assigned once in each category. This is especially problematic for Amoebozoa with complex life cycles (Keller et al., 2022; Tice et al., 2016). Consequently, this database is a simplified approximation for amoebozoan traits and contains a sum of compromises to increase its practical application.

### 2.1 Justification of genus-level trait assignment

We consider the genus level to be most suitable for assigning functional traits, as most traits (e.g., nutrition, locomotion, morphotype) in protists are conserved at the genus level (Dumack et al., 2020). In addition, sequences in reference databases are typically not assigned to species, as short reads in environmental sequencing data often do not allow for reliable taxonomic assignment at the species level. However, some traits, particularly cell size, may differ considerably between taxa, even between species within one genus, or between the different stages in the life cycle (Berney et al., 2015; Kylin, 2001). Therefore, instead of recording size as a continuous variable, we assigned a fixed (common) size range to each genus, which, however, needs to be considered with care, as variability can be large. Accordingly, comments and references are given for each genus. Moreover, although amoebozoans are phylogenetically more divergent to Cercozoa than animals to plants, we tried to keep the traits most comparable to the already published Cercozoa database (Dumack et al., 2020) but accounted also for traits intrinsic to Amoebozoa (for instance spore formation which is absent in Cercozoa).

### 2.2 Functional traits

We considered the organisms’ prey range, rough morphology, and morphotype, locomotion, known habitat preference, animal disease-relatedness (whether as vector or immediate parasite), presence/absence of spore formation, and size range. Prey range categories were grouped according to bacterivory, omnivory (feeding on bacteria and eukaryotes), eukaryvory (feeding on fungi, microfauna, algae, or other protists), and saprotrophy. We could not assign more precise categories (e.g., fungivory, algivory) for a lack of information (or contradictory reports, i.e., likely multiple trophic modes) for most taxa. Morphology was mainly specified by the presence/absence of a shell and flagella. As amoebozoan amoebae, although variable, show well-recognizable shapes(Smirnov and Brown, 2004), we further defined simplified morphotypes, i.e., disc, tubule, palm, and reticulate. Two main locomotion modes were recognized: organisms bound to the substrate, i.e., gliding or freely swimming. However, amoebae, amoeboflagellates, and flagellates differ not only in their locomotion but amoeboid cells are surface feeders, whereas prey capture of flagellates likely is much more selective due to their larger handling time. We considered habitat preferences of soil and freshwater taxa and marine taxa because soil-inhabiting and freshwater taxa may easily switch habitats, while marine taxa are rarely found in terrestrial or limnic habitats (Smirnov and Brown, 2004). Genera accommodating species from marine and soil and freshwater habitats were considered to have evolved ubiquitous habitat preferences. Spore formation is an important trait to enhance dispersal. For simplicity, we did not consider different spore formation strategies. Suggestions for updates can be addressed to the corresponding author. All consulted references fot the trait database (Supplementary Table S1) are provided the Supplementary Data S1. An R package for the easy assignment of the traits will be available at Zenodo (DOI: https://doi.org/10.5281/zenodo.15091355), updated versions of the database will be available at https://github.com/JFreude/Functional-Traits-Amoebozoa.

### 2.3 Statistics

The statistical data analyses were conducted in R v. 4.3.1 (R Core Team, 2023). The data were visualized with the R packages ggplot2 v. 3.5.1 and ggpubr v. 0.6.0 (Kassambara, 2023a; Wickham, 2011). An overview of the relative genus richness per functional traits within each category of the Amoebozoa database was given by a Sankey diagram. Only taxa for which a trait could be assigned to the respective category were considered for the relative genus richness. To explore potential convergent evolution between terrestrial Amoebozoa and Cercozoa, we also included data for Cercozoa and Endomyxa (Rhizaria) and their respective functional trait database (Dumack et al., 2020). For convenience, we will refer to Cercozoa and Endomyxa as Cercozoa.

For comparing the sizes of bacterivorous, eukaryvorous, and omnivorous taxa, the given size was used, or the mean size was calculated if a size range was given. Specifications such as “up to” or “larger than” were not considered, taxa with a size of “up to macroscopic” were regarded as 1000 µm in size. The sizes across feeding types were compared using a Kruskal-Wallis test and Dunn’s post-hoc test (rstatix v. 0.7.2::kruskal_test and rstatix v. 0.7.2:: dunn_test (Kassambara, 2023b). Pairwise comparisons were corrected for multiple testing according to Benjamini & Hochberg (1995).

To showcase the usage of the trait database, we visualized the variations in functional traits of Amoebozoa and Cercozoa communities across different habitats. We used publicly available metatranscriptomic datasets from bark (Freudenthal et al., 2024), litter (Voss et al., 2019), and soil (Fiore-Donno et al., 2024). From the latter, we only used samples that were collected in the summer (see Fiore-Donno et al., 2024). After assigning the traits, the mean and standard deviation of the relative community composition for each trait category were calculated and visualized in a point diagram for each habitat and community (Amoebozoa and Cercozoa), respectively. The proportion of taxa with missing trait information is not shown.

## 3. Results and Discussion

We provide a functional database for Amoebozoa allowing to easily add ecological meaning to molecular studies. The database comprises functional traits, i.e., habitat, locomotion, nutrition morphology, and size (Fig. 2). Additionally, we included information on morphotype, if spore formation was observed, or if they may cause diseases.

**Figure 2:**
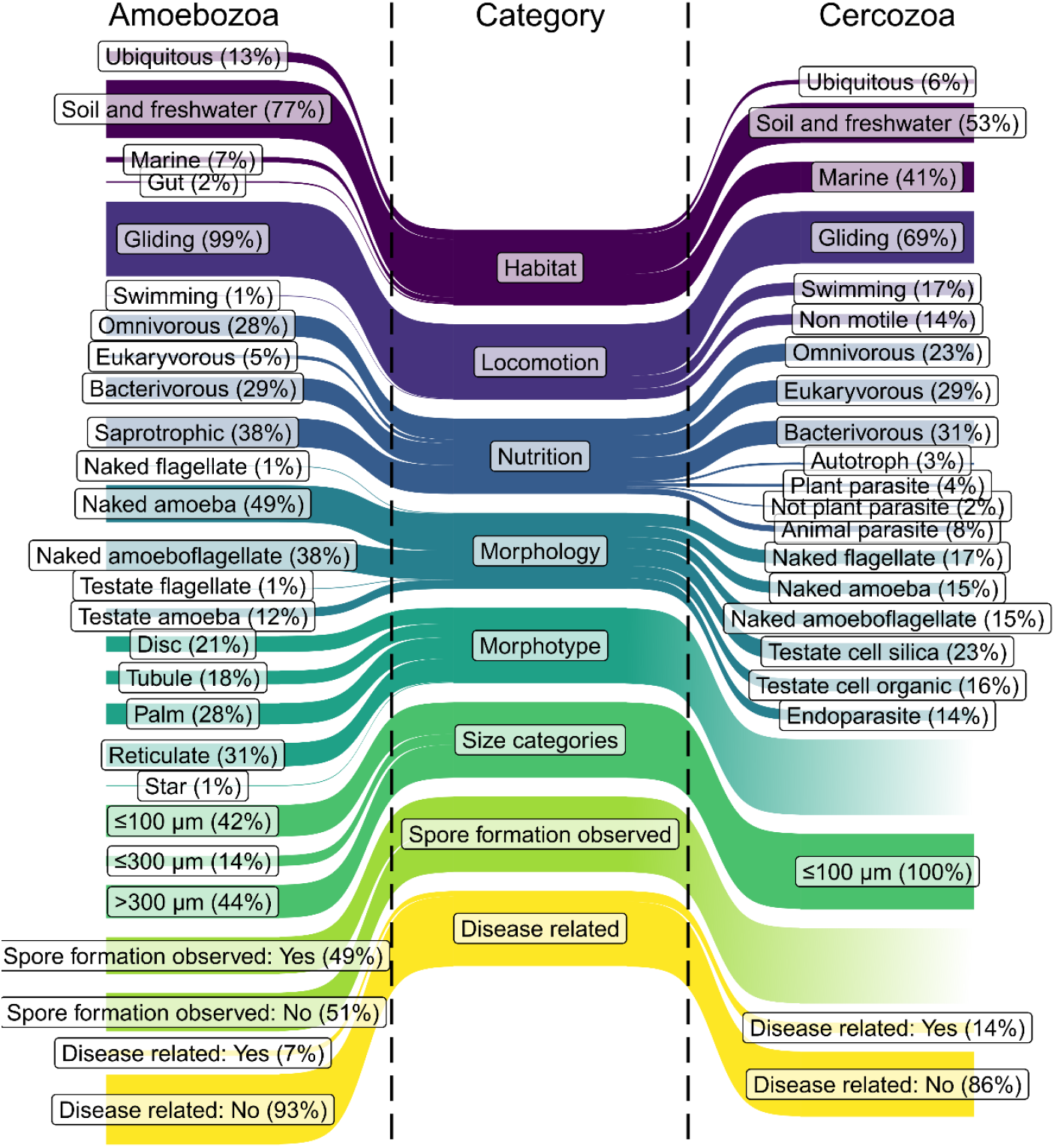
Overview of the relative genus richness per functional trait within each category of the Amobozoa and Cercozoa databases. The Sankey diagrams show the percentual genus richness calculated for the given traits of each category for the Amoebozoa (left) and Cercozoa (right) databases.

A comparison of the genus richness per trait of Amoebozoa with the Cercozoa revealed striking similarities but also distinct differences (Fig. 2). Both taxa show convergent evolution, i.e., both groups show similar morphological variation as they include shell-bearing amoebae, naked amoebae, and flagellated taxa. Furthermore, the majority of species are gliding, an adaptation to surface feeding in soil habitats. However, next to similarities, our database revealed striking differences. For example, almost 80 % of the known amoebozoan genera occur in soil or freshwater and only a small fraction in marine environments, whereas for Cercozoa, the ratio of soil and freshwater to marine genera is nearly balanced.

The mean size of Amobozoa and Cercozoa was associated with the feeding type, i.e., bacterivorous taxa were significantly smaller compared to eukaryvorous and omnivorous taxa (Fig. 3). Traditionally, most protists were considered to be bacterivorous. In recent years, however, it has become increasingly clear that many protists indeed exhibit a broad prey spectrum, including algae, fungi, and other heterotrophic protists (Dumack et al., 2019b; Estermann et al., 2023; Geisen et al., 2016; Seppey et al., 2017). For testate amoebae, it has been shown that feeding type can be determined by shell sizes, although with certain limitations (Dumack et al., 2024; Fournier et al., 2015). We now show that this also applies much more broadly to the entirety of Amobozoa and Cercozoa. Moreover, Amoebozoa comprise only a very limited number of eukaryvorous taxa, in particular in contrast to Cercozoa. While small Amoebozoa ingest only bacteria, larger ones consume bacteria and single-celled and multicellular eukaryotes., such as nematodes and fungi (Geisen et al., 2015). In other words, if the prey item can be entirely enclosed by an amoebozoan cell, it is suitable prey. Thus, the Amoebozoa are likely much less selective in their prey choice than Cercozoa, but the size of the amoebozoan cell determines which prey can be ingested (Kulishkin et al., 2023). However, few reliable data on feeding preferences exist for Amoebozoa, and more feeding experiments are urgently needed, predominantly in taxa where individual species may differ by several orders of magnitude in cell size, for example, in Variosea (Berney et al., 2015). Nonetheless, aside this generalized pattern that we found, there are some highly specialized consumers among Amoebozoa (Dumack et al., 2024, 2019a; Estermann et al., 2023; Smirnov et al., 2011), such as *Phryganella paradoxa* (Arcellinida) which feeds on pennate diatoms by bending or breaking their frustules (Dumack et al., 2024). However, these specialized consumers appear to be more exceptions than the rule.

**Figure 3:**
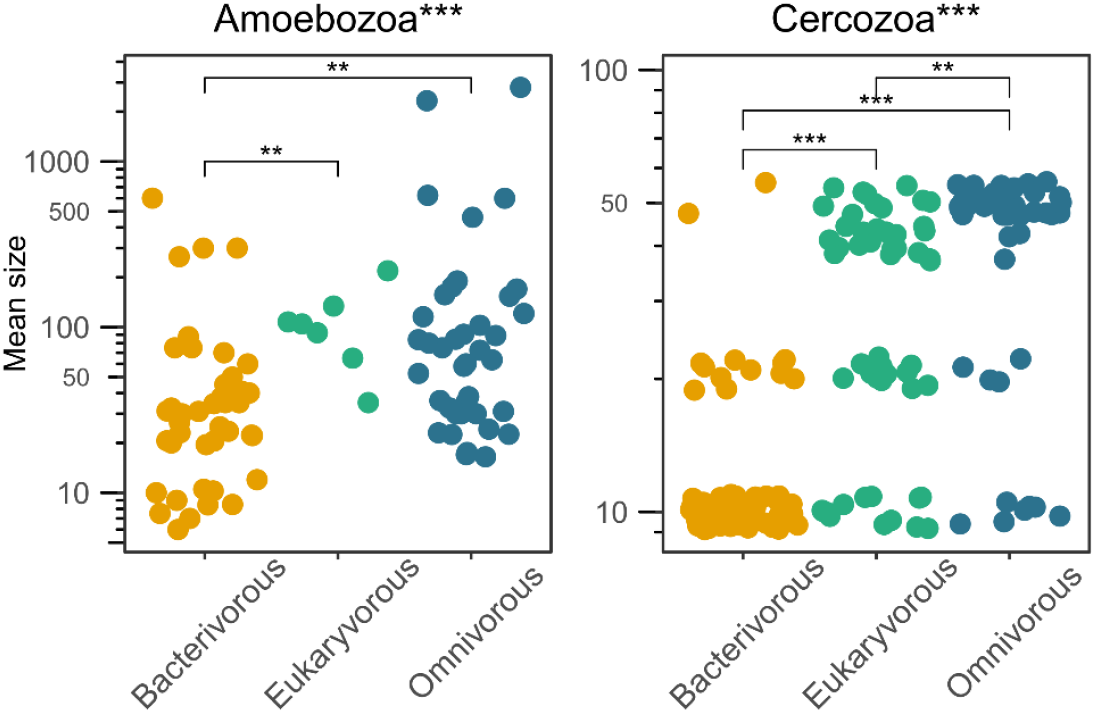
Overview of the association between size and feeding type for Amoebozoa and Cercozoa. The point diagrams show the mean sizes of bacterivorous (yellow), eukaryvorous (green) and omnivorous (blue) for the Amoebozoa (left) and Cercozoa (right) databases. Significant differences across all feeding types (Kruskal-Wallis test) and of pairwise comparisons of the feeding types (Dunn’s test) are indicated with stars in the graph title or the graph, respectively (* p < 0.05; ** p < 0.01; *** p < 0.001).

To illustrate the assignment and analyses of the Amoebozoa and Cercozoa traits, we applied the databases on recently published metatranscriptomic data sets of soil, leaf litter, and bark surfaces (Fig. 4). These datasets originate from different locations and years, providing independent inventories for the comparison of traits among communities. For example, the proportion of disease-related amoebozoa was higher for soil (∼20 %) than for bark and litter. Furthermore, spore-forming (∼90%) and saprotrophic (∼30%) Amoebozoa were exceptionally dominant on bark. Communities of Amoebozoa and terrestrial Cercozoa differed in trait composition: Omnivorous Cercozoa were most dominant on bark, while omnivorous Amoebozoa were the least abundant. In addition, the variation of functional traits between bark, litter, and soil was much lower in Cercozoa communities than within Amoebozoa.

**Figure 4:**
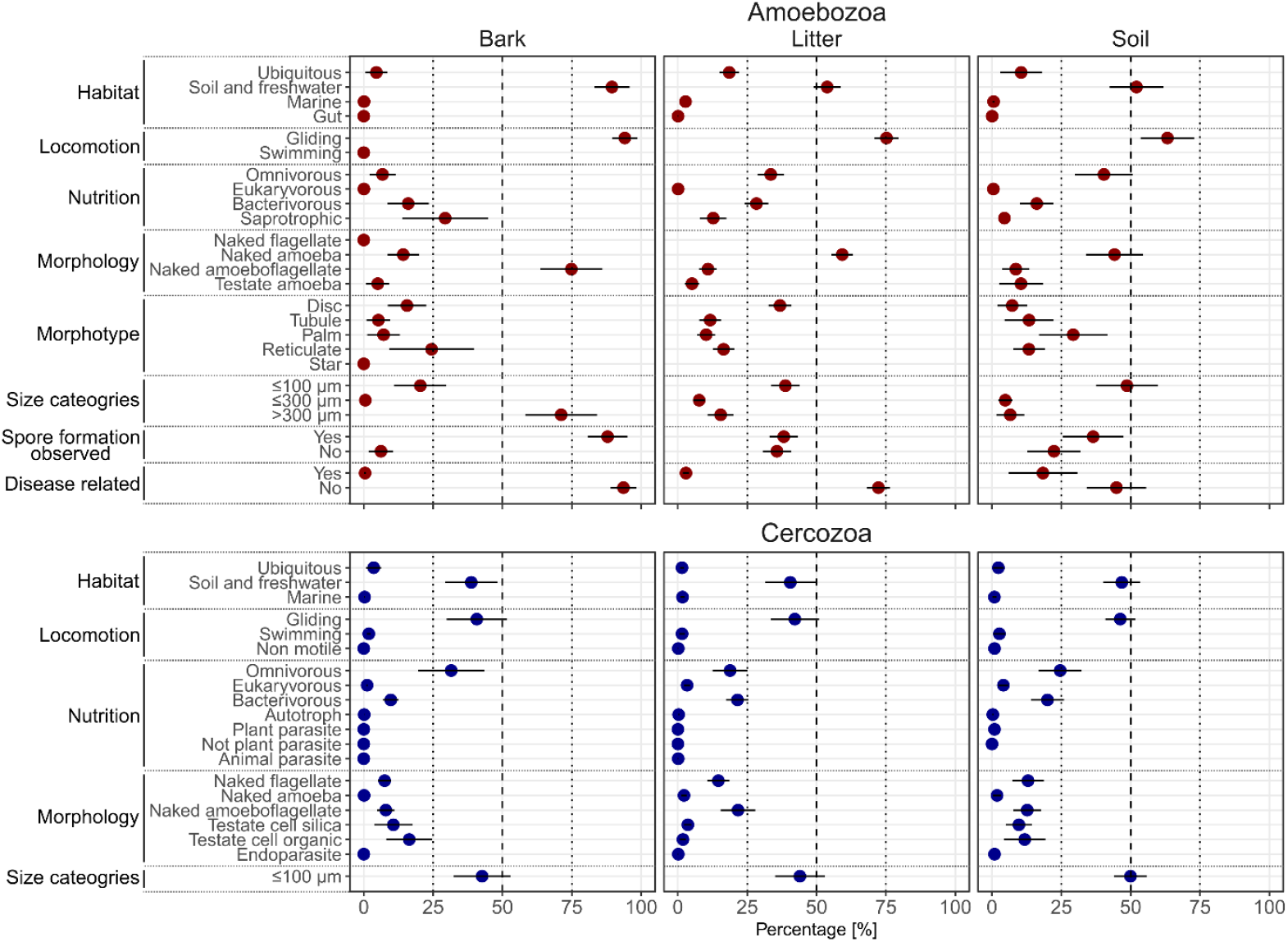
The relative proportion of functional traits of Amoebozoa and Cercozoa communities of bark, litter and soil. The point diagrams show the percentages that were assigned to the given traits of each category for Amoebozoa and Cercozoa communities of bark (N=15), litter (N=18) and soil (N=39), respectively. The points represent the mean and are color-coded by Amoebozoa (red) and Cercozoa (blue). The error bars represent the standard deviation.

The newly provided trait database for Amoebozoa enables an easy assignment of traits to environmental sequencing surveys. This will allow to detect trade-offs and evolutionary trajectories in adaptations among different supergroups in protists, and deepen our knowledge of the functional diversity of Amoebozoa and their impact on ecosystem functioning.

## 5 Data Accessibility

### Data Accessibility

The trait database and an R package for the automatic assignment of the traits to a taxonomy table will be publicly accessible upon publication at GitHub (https://github.com/JFreude/FunctionalTraitsAmoebozoa) and Zenodo (DOI: https://doi.org/10.5281/zenodo.15091355).

## 7 Acknowledgements

We would like to thank Ferry Siemensma for providing the amazing pictures of the Amoebozoa.

## 8 Author Contributions

KD, MS and MB designed the study. KD and JF compiled the trait database. JF created all figures, performed the statistical analyses and wrote the R package. JF outlined the manuscript, all co-authors commented on the manuscript and approved the submitted version.

