## Supplementary Data S1 for "A novel protistan trait database reveals functional redundancy and complementarity in terrestrial protists (Amoebozoa & Rhizaria)"

***Functional traits of Amoebozoa, references used in Table S1.***


Angell, R.W., 1976. Observations on Trichosphaerium platyxyrum sp. n. J. Protozool. 23, 357–364. https://doi.org/10.1111/j.1550-7408.1976.tb03788.x

Baumgartner, M., Yapi, A., Gröbner-Ferreira, R., Stetter, K.O., 2003. Cultivation and properties of *Echinamoeba thermarum* n. sp., an extremely thermophilic amoeba thriving in hot springs. Extremophiles 7, 267–274. https://doi.org/10.1007/s00792-003-0319-6

Berney, C., Geisen, S., Van Wichelen, J., Nitsche, F., Vanormelingen, P., Bonkowski, M., Bass, D., 2015. Expansion of the 'Reticulosphere': Diversity of novel branching and network-forming amoebae helps to define Variosea (Amoebozoa). Protist 166, 271–295. https://doi.org/10.1016/j.protis.2015.04.001

Blandenier, Q., Seppey, C.V.W., Singer, D., Vlimant, M., Simon, A., Duckert, C., Lara, E., 2017. *Mycamoeba gemmipara* nov. gen., nov. sp., the First Cultured Member of the Environmental Dermamoebidae Clade LKM74 and its Unusual Life Cycle. J. Eukaryot. Microbiol. 64, 257–265. https://doi.org/10.1111/jeu.12357

Bobrov, A., Kosakyan, A., 2015. A New Species from Mountain Forest Soils in Japan: *Porosia paracarinata* sp. nov., and Taxonomic Concept of the Genus *Porosia* Jung, 1942. Acta Protozool. 2015, 289–294. https://doi.org/10.4467/16890027AP.15.024.3538

Bobrov, A., Mazei, Y., 2017. A review of testate amoeba genus *Cryptodifflugia* Penard, 1890 (Phryganellina: Cryptodifflugiidae) with a key to species. Zootaxa 4282, 292–308. https://doi.org/10.11646/zootaxa.4282.2.4

De Saedeleer, H., 1934. Beitrag zur Kenntnis der Rhizopoden: morphologische und systematische Untersuchungen und ein Klassifikationsversuch. Mem. R. Belg. Mus. Nat. Sci. 60, 3–112.

Dumack, K., Kahlich, C., Lahr, D.J.G., Bonkowski, M., 2019. Reinvestigation of *Phryganella paradoxa* (Arcellinida, Amoebozoa) Penard 1902. J. Eukaryot. Microbiol. 66, 232–243. https://doi.org/10.1111/jeu.12665

Dyková, I., Kostka, M., Pecková, H., 2010. *Grellamoeba robusta* gen. n., sp. n., a possible member of the family Acramoebidae Smirnov, Nassonova et Cavalier-Smith, 2008. Eur. J. Protistol. 46, 77–85. https://doi.org/10.1016/j.ejop.2009.10.004

Dyková, I., Veverková-Fialová, M., Fiala, I., Dvořáková, H., 2005. *Protacanthamoeba bohemica* sp. n., Isolated from the Liver of Tench, *Tinca tinca* (Linnaeus, 1758). Acta Protozool.

El-Dib, N.A., 2017. *Entamoeba histolytica*: an Overview. Curr. Trop. Med. Rep. 4, 11–20. https://doi.org/10.1007/s40475-017-0100-z

Geisen, S., Weinert, J., Kudryavtsev, A., Glotova, A., Bonkowski, M., Smirnov, A., 2014. Two new species of the genus *Stenamoeba* (Discosea, Longamoebia): Cytoplasmic MTOC is present in one more amoebae lineage. Eur. J. Protistol. 50, 153–165. https://doi.org/10.1016/j.ejop.2014.01.007

Goodkov, A.V., 1988. Korotnevella nom. nov.—new generic name for scale-bearing Mayorella-like amoebae. Zool. Zhurnal 67, 1728–1730.

Iwamoto, Y., Degawa, Y., Nakayama, T., 2023. Re-examination of a rare protosteloid amoeba *Schizoplasmodiopsis micropunctata*, and the revision of *Tychosporium* (Cavosteliida, Variosea, Amoebozoa). Mycoscience 64, 63–68. https://doi.org/10.47371/mycosci.2023.01.002

Kang, S., Tice, A.K., Spiegel, F.W., Silberman, J.D., Pánek, T., Čepička, I., Kostka, M., Kosakyan, A., Alcântara, D.M.C., Roger, A.J., Shadwick, L.L., Smirnov, A., Kudryavtsev, A., Lahr, D.J.G., Brown, M.W., 2017. Between a Pod and a Hard Test: The Deep Evolution of Amoebae. Mol. Biol. Evol. 34, 2258–2270. https://doi.org/10.1093/molbev/msx162

Kosakyan, A., Heger, T.J., Leander, B.S., Todorov, M., Mitchell, E.A.D., Lara, E., 2012. COI Barcoding of Nebelid Testate Amoebae (Amoebozoa: Arcellinida): Extensive Cryptic Diversity and Redefinition of the Hyalospheniidae Schultze. Protist 163, 415–434. https://doi.org/10.1016/j.protis.2011.10.003

Kosakyan, A., Meisterfeld, R., Lara, E., Duckert, C., Mitchell, E.A.D., 2025. A taxonomic monograph of hyalospheniid testate amoebae (Amoebozoa: Arcellinida: Hyalospheniformes), 1st ed. Éditions Alphil-Presses universitaires suisses. https://doi.org/10.33055/ALPHIL.00614
Kudryavtsev, A., 2006. “Minute” species of *Cochliopodium* (Himatismenida): Description of three new fresh- and brackish-water species with a new diagnosis for *Cochliopodium minus* Page, 1976. Eur. J. Protistol. 42, 77–89. https://doi.org/10.1016/j.ejop.2005.12.002
Kudryavtsev, A., Hausmann, K., 2007. *Spumochlamys iliensis* n.g. n. sp. (Testacealobosia, Microchlamyiidae) from Central Asia, with notes on the diversity of *Microchlamys*-like testate amoebae. Eur. J. Protistol. 43, 185–191. https://doi.org/10.1016/j.ejop.2007.03.001
Kudryavtsev, A., Pawlowski, J., 2015. *Cunea* n. g. (Amoebozoa, Dactylopodida) with two cryptic species isolated from different areas of the ocean. Eur. J. Protistol. 51, 197–209. https://doi.org/10.1016/j.ejop.2015.04.002
Kudryavtsev, A., Pawlowski, J., 2013. *Squamamoeba japonica* n. g. n. sp. (Amoebozoa): A Deep-sea Amoeba from the Sea of Japan with a Novel Cell Coat Structure. Protist 164, 13–23. https://doi.org/10.1016/j.protis.2012.07.003
Kudryavtsev, A., Völcker, E., Clauß, S., Pawlowski, J., 2021. *Ovalopodium rosalinum* sp. nov., *Planopodium haveli* gen. nov, sp. nov., *Planopodium desertum* comb. nov. and new insights into phylogeny of the deeply branching members of the order Himatismenida (Amoebozoa). Int. J. Syst. Evol. Microbiol. 71, 004737. https://doi.org/10.1099/ijsem.0.004737
Kudryavtsev, A., Volkova, E., 2018. *Clydonella sawyeri* n. sp. (Amoebozoa, Vannellida): Morphological and molecular study and a re-definition of the genus *Clydonella* Sawyer, 1975. Eur. J. Protistol. 63, 62–71. https://doi.org/10.1016/j.ejop.2018.01.008
Kudryavtsev, A., Wylezich, C., Schlegel, M., Walochnik, J., Michel, R., 2009. Ultrastructure, SSU rRNA Gene Sequences and Phylogenetic Relationships of *Flamella* Schaeffer, 1926 (Amoebozoa), with Description of Three New Species. Protist 160, 21–40. https://doi.org/10.1016/j.protis.2008.09.004
Lado, C., Basanta, D.W. de, 2008. A Review of Neotropical Myxomycetes (1828-2008). An. Jardín Botánico Madr. 65, 211–254. https://doi.org/10.3989/ajbm.2008.v65.i2.293
Leidy, J., 1879. Fresh-water rhizopods of North America. Government Printing Office, Washington.
Mikrjukov, K.A., Mylnikov, A.P., 1998. The fine structure of a carnivorous multiflagellar protist *Multicilia marina* Cienkowski, 1881 (flagellata incertae sedis). Eur. J. Protistol. 34, 391–401. https://doi.org/10.1016/S0932-4739(98)80008-4
Ntakou, E., Siemensma, F., Bonkowski, M., Dumack, K., 2019. The Dancing Star: Reinvestigation of *Artodiscus saltans* (Variosea, Amoebozoa) Penard 1890. Protist 170, 349–357. https://doi.org/10.1016/j.protis.2019.06.002
Olive, L.S., Bennett, W.E., Deasey, M.C., 1984. The New Protostelid Genus *Endostelium*. Mycologia 76, 884–891. https://doi.org/10.2307/3793145
Olive, L.S., Stoianovich, C., 1972. *Protosporangium*: a New Genus of Protostelids. J. Protozool. 19, 563–571. https://doi.org/10.1111/j.1550-7408.1972.tb03530.x
Olive, L.S., Stoianovitch, C., 1979. Observations on the Mycetozoan Genus *Ceratiomyxa*: Description of a New Species. Mycologia 71, 546–555. https://doi.org/10.2307/3759064
Olive, L.S., Stoianovitch, C., 1977. *Clastostelium*, a new ballistosporous protostelid (Mycetozoa) with flagellate cells. Trans. Br. Mycol. Soc. 69, 83–88. https://doi.org/10.1016/S0007-1536(77)80119-8
Olive, L.S., Stoianovitch, C., 1975. The Protostelid Genus *Schizoplasmodiopsis*. Mycologia 67, 1087–1100. https://doi.org/10.1080/00275514.1975.12019851
Olive, L.S., Stoianovitch, C., 1971a. A New Genus of Protostelids Showing Affinities with *Ceratiomyxa*. Am. J. Bot. 58, 32–40. https://doi.org/10.1002/j.1537-2197.1971.tb09942.x
Olive, L.S., Stoianovitch, C., 1971b. *Planoprotostelium*, a New Genus of Protostelids. J. Elisha Mitchell Sci. Soc. 87, 115–119.
Olive, L.S., Stoianovitch, C., 1966. A New Two-Spored Species of *Cavostelium* (Protostelida). Mycologia 58, 440–451. https://doi.org/10.1080/00275514.1966.12018335

Page, F.C., 1987. The Classification of ‘Naked’ Amoebae (Phylum Rhizopoda). Arch. Für Protistenkd. 133, 199–217. https://doi.org/10.1016/S0003-9365(87)80053-2

Page, F.C., 1980. A light-and electron-microscopical comparison of marine Umax and flabellate amoebae belonging to four genera. Protistologica 16, 57–78.

Page, F.C., 1979. Two genera of marine amoebae (Gymnamoebia) with distinctive surface structures: *Vannella* Bovee, 1965, and *Pseudoparamoeba* n. gen., with two new species of *Vannella*. Protistologica 15, 243–255.

Page, F.C., 1976. A revised classification of the Gymnamoebia (Protozoa: Sarcodina). Zool. J. Linn. Soc. 58, 61–77. https://doi.org/10.1111/j.1096-3642.1976.tb00820.x

Page, F.C., 1972. A study of two *Mayorella* species and proposed union of the families Mayorellidae and Paramoebidae (Rhizopodea, Amoebida). Arch. Für Protistenkd. 114, 404–420.

Page, F.C., Blakey, S.M., 1979. Cell surface structure as a taxonomic character in the Thecamoebidae (Protozoa: Gymnamoebia). Zool. J. Linn. Soc. 66, 113–135. https://doi.org/10.1111/j.1096-3642.1979.tb01905.x

Penard, E., 1907. On Some Rhizopods from the Sikkim Himalaya. J. R. Microsc. Soc. 27, 274–278.

Polne-Fuller, M., Rogerson, A., Amano, H., Gibor, A., 1990. Digestion of seaweeds by the marine amoeba *Trichosphaerium*. Hydrobiologia 204, 409–413. https://doi.org/10.1007/BF00040264

Poulsen, C.S., Stensvold, C.R., 2016. Systematic review on *Endolimax nana*: A less well studied intestinal ameba. Trop. Parasitol. 6, 8. https://doi.org/10.4103/2229-5070.175077

Ptáčková, E., Kostygov, A.Yu., Chistyakova, L.V., Falteisek, L., Frolov, A.O., Patterson, D.J., Walker, G., Cepicka, I., 2013. Evolution of Archamoebae: Morphological and Molecular Evidence for Pelobionts Including *Rhizomastix*, *Entamoeba*, *Iodamoeba*, and *Endolimax*. Protist 164, 380–410. https://doi.org/10.1016/j.protis.2012.11.005

Radosa, S., Ferling, I., Sprague, J.L., Westermann, M., Hillmann, F., 2019. The different morphologies of yeast and filamentous fungi trigger distinct killing and feeding mechanisms in a fungivorous amoeba. Environ. Microbiol. 21, 1809–1820. https://doi.org/10.1111/1462-2920.14588

Rogerson, A., 1993. *Parvamoeba rugata* n. g., n. sp., (Gymnamoebia, Thecamoebidae): An exceptionally small marine naked amoeba. Eur. J. Protistol. 29, 446–452. https://doi.org/10.1016/S0932-4739(11)80407-4

Sawyer, T.K., 1975. Marine Amoebae from Surface Waters of Chincoteague Bay, Virginia: One New Genus and Eleven New Species within the Families Thecamoebidae and Hyalodiscidae. Trans. Am. Microsc. Soc. 94, 305–323. https://doi.org/10.2307/3225496

Schaeffer, A.A., 1916. Notes on the specific and other characters of Amoeba proteus Pallas (Leidy), *A. discoides* spec. nov., and *A. dubia* spec. nov. Arch. Für Protistenkd. 37, 204–228.

Schaudinn, F.R., 1896. Über den Zeugungskreis von *Paramoeba eilhardi* n.g. n.sp. Sitzungsberichte K. Preuss. Akad. Wiss. Zu Berl. 14, 31–41.

Siddiqui, R., Makhlouf, Z., Khan, N.A., 2021. The increasing importance of *Vermamoeba vermiformis*. J. Eukaryot. Microbiol. 68, e12857. https://doi.org/10.1111/jeu.12857

Siemensma, F., n.d. Microworld – world of amoeboid organisms. URL https://arcella.nl/, https://arcella.nl/ (accessed 3.27.25).

Smirnov, A., Nassonova, E., Fahrni, J., Pawlowski, J., 2009. *Rhizamoeba neglecta* n. sp. (Amoebozoa, Tubulinea) from the bottom sediments of freshwater Lake Leshevoe (Valamo Island, North-Western Russia), with notes on the phylogeny of the order Leptomyxida. Eur. J. Protistol. 45, 251–259. https://doi.org/10.1016/j.ejop.2009.04.002

Smirnov, A., Nassonova, E., Geisen, S., Bonkowski, M., Kudryavtsev, A., Berney, C., Glotova, A., Bondarenko, N., Dyková, I., Mrva, M., Fahrni, J., Pawlowski, J., 2017. Phylogeny and Systematics of Leptomyxid Amoebae (Amoebozoa, Tubulinea, Leptomyxida). Protist 168, 220–252. https://doi.org/10.1016/j.protis.2016.10.006

Smirnov, A.V., 1997. Two new species of marine amoebae: *Hartmannella lobifera* n. sp. and *Korotnevella nivo* n. sp. (Lobosea, Gymnamoebida). Arch. Für Protistenkd. 147, 283–292. https://doi.org/10.1016/S0003-9365(97)80055-3

Smirnov, A.V., Goodkov, A.V., 1993. *Paradermamoeba valamo* gen. n., sp. n. (Gymnamoebia, Thecamoebidae) - a freshwater amoeba from bottom sediments. Zool. Zhurnal 72, 5–11.
Smirnov, A.V., Kudryavtsev, A.A., 2005. Pellitidae n. fam. (Lobosea, Gymnamoebia) – a new family, accommodating two amoebae with an unusual cell coat and an original mode of locomotion, *Pellita catalonica* n.g., n.sp. and *Pellita digitata* comb. nov. Eur. J. Protistol. 41, 257–267. https://doi.org/10.1016/j.ejop.2005.05.002
Smirnov, A.V., Nassonova, E.S., Cavalier-Smith, T., 2008. Correct identification of species makes the amoebozoan rRNA tree congruent with morphology for the order Leptomyxida Page 1987; with description of *Acramoeba dendroida* n. g., n. sp., originally misidentified as '*Gephyramoeba* sp.' Eur. J. Protistol. 44, 35–44. https://doi.org/10.1016/j.ejop.2007.08.001
Smirnov, A.V., Nassonova, E.S., Chao, E., Cavalier-Smith, T., 2007. Phylogeny, Evolution, and Taxonomy of Vannellid Amoebae. Protist 158, 295–324. https://doi.org/10.1016/j.protis.2007.04.004
Spiegel, F.W., Gecks, S.C., Feldman, J., 1994. Revision of the Genus *Protostelium* (Eumycetozoa) I: The *Protostelium* mycophaga Group and the *P. irregularis* Group. J. Eukaryot. Microbiol. 41, 511–515. https://doi.org/10.1111/j.1550-7408.1994.tb06051.x
Stephenson, S.L., Fiore-Donno, A.M., Schnittler, M., 2011. Myxomycetes in soil. Soil Biol. Biochem. 43, 2237–2242. https://doi.org/10.1016/j.soilbio.2011.07.007
Tice, A.K., Shadwick, L.L., Fiore-Donno, A.M., Geisen, S., Kang, S., Schuler, G.A., Spiegel, F.W., Wilkinson, K.A., Bonkowski, M., Dumack, K., Lahr, D.J.G., Voelcker, E., Clauß, S., Zhang, J., Brown, M.W., 2016. Expansion of the molecular and morphological diversity of Acanthamoebidae (Centramoebida, Amoebozoa) and identification of a novel life cycle type within the group. Biol. Direct 11, 69. https://doi.org/10.1186/s13062-016-0171-0
Tyml, T., Kostka, M., Ditrich, O., Dyková, I., 2016. *Vermistella arctica* n. sp. Nominates the Genus *Vermistella* as a Candidate for Taxon with Bipolar Distribution. J. Eukaryot. Microbiol. 63, 210–219. https://doi.org/10.1111/jeu.12270
Van Wichelen, J., D 'Hondt, S., Claeys, M., Vyverman, W., Berney, C., Bass, D., Vanormelingen, P., 2016. A Hotspot of Amoebae Diversity: 8 New Naked Amoebae Associated with the Planktonic Bloom-forming Cyanobacterium *Microcystis*. Acta Protozool. 55, 61. https://doi.org/10.4467/16890027AP.16.007.4942
Visvesvara, G.S., Moura, H., Schuster, F.L., 2007. Pathogenic and opportunistic free-living amoebae: *Acanthamoeba* spp., *Balamuthia mandrillaris*, *Naegleria fowleri*, and *Sappinia diploidea*. FEMS Immunol. Med. Microbiol. 50, 1–26. https://doi.org/10.1111/j.1574-695X.2007.00232.x
Watson, P.M., Sorrell, S.C., Brown, M.W., 2014. *Ptolemeba* n. gen., a Novel Genus of Hartmannellid Amoebae (Tubulinea, Amoebozoa); with an Emphasis on the Taxonomy of *Saccamoeba*. J. Eukaryot. Microbiol. 61, 611–619. https://doi.org/10.1111/jeu.12139
Zadrobílková, E., Walker, G., Čepička, I., 2015. Morphological and Molecular Evidence Support a Close Relationship Between the Free-living Archamoebae *Mastigella* and *Pelomyxa*. Protist 166, 14–41. https://doi.org/10.1016/j.protis.2014.11.003
